# Human primary motor cortex indexes the onset of subjective intention in brain-machine-interface mediated actions

**DOI:** 10.1101/2023.07.21.550067

**Authors:** Jean-Paul Noel, Marcia Bockbrader, Sam Colachis, Marco Solca, Pavo Orepic, Patrick D. Ganzer, Patrick Haggard, Ali Rezai, Olaf Blanke, Andrea Serino

**Author notes:** These authors contributed equally (Jean-Paul Noel, Marcia Bockbrader, Olaf Blanke, Andrea Serino). These authors jointly supervised this work and are listed in alphabetic order.

## Abstract

Self-initiated behavior is accompanied by the experience of willing our actions. Here, we leverage the unique opportunity to examine the full intentional chain – from will (W) to action (A) to environmental effects (E) - in a tetraplegic person fitted with a primary motor cortex (M1) brain machine interface (BMI) generating hand movements via neuromuscular electrical stimulation (NMES). This combined BMI-NMES approach allowed us to selectively manipulate each element of the intentional chain (W, A, and E) while performing extra-cellular recordings and probing subjective experience. Our results reveal single-cell, multi-unit, and population-level dynamics in human M1 that encode W and may predict its subjective onset. Further, we show that the proficiency of a neural decoder in M1 reflects the degree of W-A binding, tracking the participant’s subjective experience of intention in (near) real time. These results point to M1 as a critical node in forming the subjective experience of intention and demonstrate the relevance of intention-related signals for translational neuroprosthetics.

## Introduction

Self-initiated behavior is accompanied by the experience of willing our actions (1). This same subjective experience can be elicited by direct stimulation of frontal and parietal areas, with subjects reporting the “urge to move” (2, 3). Further, spiking activity from neurons in frontal areas anticipate the subjective experience of willing movements by up to ∼1.5 seconds (4). These findings suggest that a fronto-parietal circuit initiates movements *prior* to the agent being aware of this intention (5–10), fueling a long-term debate regarding the nature of free-will. However, the neural mechanisms underlying (i.e., co-occuring with) the subjective experience of intending actions remain unknown.

The study of intention has largely relied on two paradigms, the “Libet task” (5) and “Intentional Binding” (11, 12). In the former, participants self-initiate movements and report the perceived time of their intention while neurophysiological measures are taken. Libet’s seminal observation (5) was that a slow drift of neural ensemble activity, the readiness potential (13), preceded not only the onset of voluntary action (A-time), but also the subject’s perceived intent to move (W-time, but see (14, 15)). In the latter (11, 12), participants perform a simple action leading to an effect on the environment. Then, participants report on the timing of their action or the consequent environmental effect. The seminal observation (11, 12) is that voluntary actions (but not involuntary ones) lead to a compression in the perceived timing between actions and effects.

Studies employing these paradigms have significantly advanced our understanding of motor intention, and have highlighted a common neural circuitry including the anterior cingulate cortex (ACC), supplementary motor areas (pre-SMA, and SMA proper), and the posterior parietal cortex, leading to self-initiated behavior (5, 11, 12, 16–21). However, these studies have not examined the full intentional chain; from will (W) to action (A) to effect (E). The voluntary actions in the “Libet Task” omit E, while “Intentional Binding” invokes W, but does not measure it. More vexingly, there is a dearth of cellular recordings on intentional processes. For instance, “Intentional Binding” has never been examined during invasive, single-cell recordings. In turn, the relative timing between the subjective experience of W, A, and E, and associated neural activity remains unclear. Finally, most previous work has focused on preparatory or precursor activity in higher-order areas of the frontal motor hierarchy, yet have failed to consider the contribution of primary motor cortex (M1) in the intentional chain. M1 is the last cortical node guiding action - what Sherrington (22) called “the final common path” - and the node most targeted in building invasive brain machine interfaces (BMIs) to restore motor control. Yet, we do not know if and how M1 reflects subjective intention, and whether such signals influence BMI performance.

To bridge these gaps, here we leveraged the unique opportunity to examine the full intentional chain in an expert intracortical M1 BMI user who is outfitted with Neuromuscular Electrical Stimulation (NMES) leading to movements of his own body (23–25). This unique setup allowed us to realize a novel intentional chain paradigm in which we systematically enable or disable each element of the intentional chain (W, A, E), while recording neural activity in M1 and collecting subjective reports regarding the perceived timing of W, A, and E.

## Results

### Binding between intention and action

The participant was a C5/C6 tetraplegic person outfitted with a Utah microelectrode array (96 electrodes) in the hand region of M1 (**Fig. 1A**). He is an expert BMI user (23–25) able to perform dexterous movements with his real forearm/hand, given that appropriate NMES is provided based on neural decoding. To this end, prior to each session the participant was asked to attempt two movements, right hand opening (HO) and closing (HC). From these, a non-linear Support Vector Machine (SVM, 26) was trained to detect intention to move (accuracy of 89.2% for HC, no false positives; see **Fig. S1** and *Methods*). During experimental sessions (total of 12 sessions, ∼35 hours total), the participant was asked to initiate HCs at a time of his own will (W). His intention to move was decoded via the BMI system and NMES was used to realize the action (A). The participant held a ball in his hand, serving as an apparent actuator causing an auditory tone 300ms after squeezing it, i.e., an effect in the environment (E). In different blocks, distinct elements of the intentional chain were bypassed by generating an involuntary hand movement via NMES (i.e., absence of W), by not activating the NMES when an intention was decoded (i.e., absence of A), or by not generating the sound after HC (i.e., absence of E). During the task, the participant viewed a rendering of a clock with a single hand that would complete a full cycle in 2560ms. On different trials, he was prompted to report the timing of a single element of the intentional chain.

**Figure 1.**
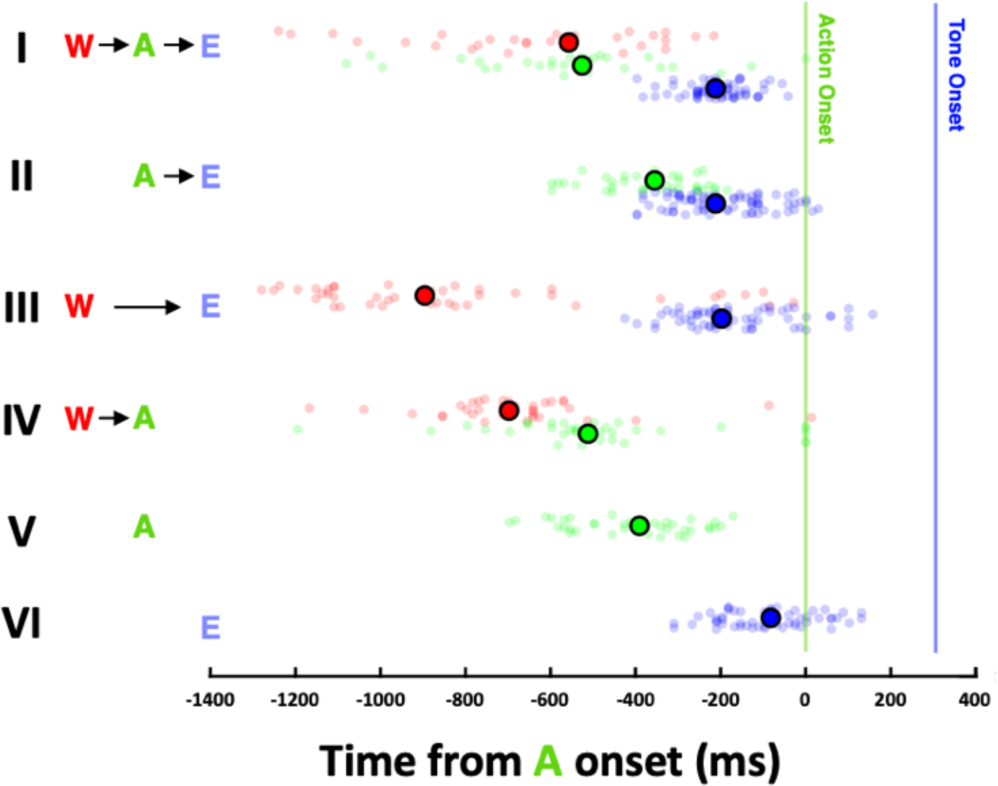
Behavioral Results. I. Full intentional chain where the BMI user indicates time of will (W, red), action (A, green), and an effect (E, blue) in the external environment. Overall, timing is biased (vertical lines are objective timing) but relative timing is accurate and precise. II. Estimates of the timing of actions and effects in the absence of intention. III. Estimates of the timing of intentions and effects in the absence of actions. IV. Estimates of the timing of intentions and actions in the absence of an effect. V. Estimate of the timing of actions in the absence of intentions and effects. VI. Estimate of the timing of effects in the absence of intentions and actions. Big circles outlined in black are median estimates, while smaller semi-transparent circles are individual trials.

Overall, the BMI participant had a bias in the temporal perception of his actions and their consequences, estimating these to occur ∼450-500ms prior to their objective timing (mean ± S.E.M.; A-Time: −455ms ± 34ms; E-Time; −512ms ± 7ms, **Fig. 1B**). More importantly, the relative timing between hand movement and the following auditory cue was accurate (median of all possible permutations of differences between A to E estimates = 257ms; objective timing difference = 300ms) and precise (S.E.M across A and E judgment types = 15ms).

Under the full intentional chain, when W led to A which in turn led to E, the participant perceived W to lead A by 71ms, and E to follow A by 314ms (error from objective timing = 14ms. **Fig. 1B, Row I**). When removing W and instead provoking HCs at arbitrary times, A were perceived to occur much later in time (with W: −526ms ± 44ms relative to objective A-time; without W; −355ms ± 18ms, p = 9.09×10^-5^, **Fig. 1B, Row II).** When removing A by transiently disabling the NMES command, and instead activating the tone at a delay after the decoder reached threshold for motor initiation (W-E), W was perceived to occur much earlier in time (with A: −597ms ± 79ms relative to A-time; without A: −796ms ± 129ms, p = 0.02, **Fig. 1B, Row III**). In neither manipulation – selectively bypassing W or A – did the estimated timing of the tone (E) change (all p > 0.1). Lastly, eliminating E from the full intentional chain did not alter the perceived timing of W (p = 0.41) or A (p = 0.07) relative to the full intentional chain (**Fig. 1B, Row IV**). These results show a novel form of intentional binding; a compression in the subjective timing between W and A.

Whereas previous binding was observed for the association between two physical events (11, 12) (A and E; see ***Supplementary Results*** for replication of this A-E binding), here we describe a new binding between a purely internal event (W) and a physical action (A). W-A binding could have not been revealed in previous work given that only in a participant with a disconnection between brain and effector can intention and motor output be independently controlled and measured.

### M1 responses index intentionality

Multi-unit activity (MUA) were aligned to movement onset, or in the case of movement-absent W trials, to when movement onset should have occurred given BMI decoding. Results demonstrated consistent MUA in M1 during intention-only trials (peak ∼5-8Hz occuring 338ms prior to movement onset, **Fig. 2A**), as well as during NMES-activated movement (peak ∼20Hz, occuring 373ms after movement onset, **Fig. 2B**). There was no MUA response to E (**Fig. S2A**). These results demonstrate that intention signals are present in M1, that they precede action-related activity, and that they are expectedly smaller than action-related activity. Further, a direct comparison of intended vs. unintended actions confirms the presence of a gradual rise in firing rate prior to action initiation only when these were intended (**Fig. S3**).

**Figure 2.**
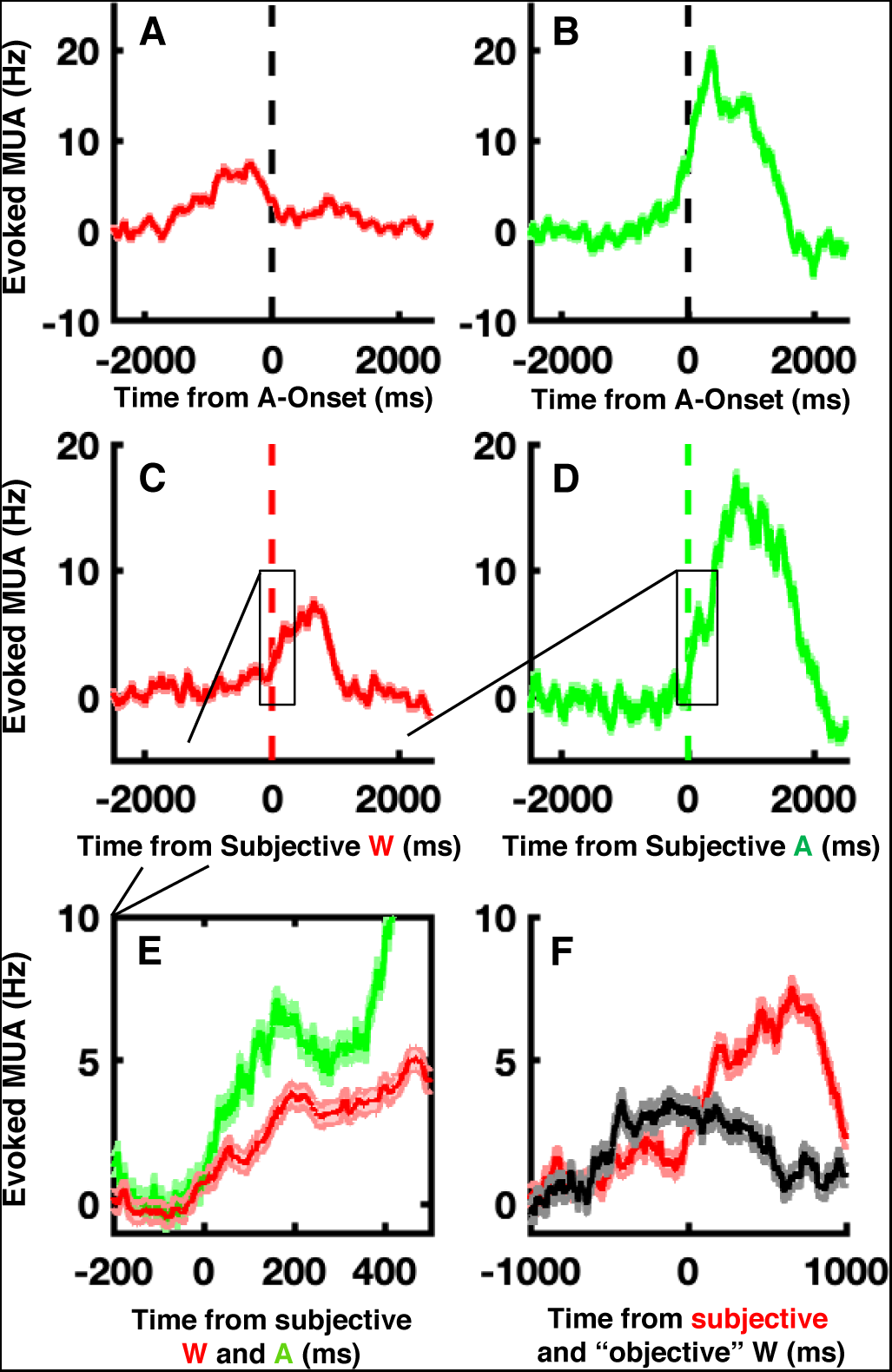
Multi-Unit Activity (MUA) in primary motor cortex (M1) for intention and action. MUA evoked during intention-only **(A,** red) and movement-only **(B,** green) conditions aligned to the objective time of movement onset (A-Onset) or decoder crossing threshold for movement. MUA evoked during intention- **(C,** red) and movement-only **(D,** green) conditions, each aligned to their subjective timing of occurrence (vertical lines in the corresponding color of intention and action **E)**. A zoomed-in version (x-axis spanning −200 to 500ms) directly contrasting MUA aligned to subjective experience during intention-only and movement-only trials. **F).** MUA during intention-only trials, aligned to the subjective timing of intention (red, same as in **C**, **E**.) and the timing at which SVM decoder was first able to detect intentionality (“objective” timing of will, black). Shaded areas surrounding the average MUA are S.E.M.

We were particularly interested in the temporal relation between this activity and the perceived timing of events along the intentional chain. Thus, we aligned MUA to the subjective timings of W, A, and E. Results showed that averaged MUA closely followed the subjective experience, being evident 14ms after the subjective estimate of W (**Fig. 2C**), and 7ms after the subjective estimate of A (**Fig. 2D**; see **Fig. 2E** for a zoomed-in version comparing W and A). No responses were found to the subjective timing of E (**Fig. S2B**). Quite uniquely, within this BMI context we could also determine on a trial-by-trial basis an “objective” timing of will. We operationalized the latter as the first time-point within a trial wherein the decoder surpassed 5 standard deviations above its noise level. Aligning MUA during no-movement trials to both “objective” and “subjective” timing of intention (**Fig. 2F**, black = “objective”, red = subjective) showed that MUA in these trials coincided with the subjective report of willing an action, but not with the “objective” timing of intention. This shows that while decoders (either artificial or downstream neural areas) may have access to an anticipatory intention signal (e.g., SMA or ACC), the experience of intention co-occurs with clear evoked spiking activity in M1.

To study population dynamics, we projected averaged spiking activity for all 96 channels onto their first two principal components. This accounted for 81.9% of the total variance. The contrast between conditions differing solely by the presence (vs. absence) of W showed that trajectories in a low dimensional latent space bifurcated (**Fig. S4A**). The distance (Euclidian in 2D) between these conditions gradually grew prior to movement, peaked 300ms post movement onset, and then precipitously dropped (**Fig. S4A**; see **Fig. S4B** and **C** for the same analyses for trials missing either A or E). We further investigated whether population dynamics in M1 reflect subjective experience by testing whether they discriminate between trials matched for the presence of W, A, and E, but differing in their subjective temporal experience. The contrast between trials where W was perceived relatively early vs. late differed between −1360ms and −360ms relative to movement onset (**Fig. S5A,** see **Fig. S5B** and **C** for A and E).

Overall, these results indicate that average MUA responses in M1 index the presence of intention and are sensitive to its perceived timing. MUA co-occurs with the onset of the experience of intention. This suggests that M1 plays an important role in eliciting the subjective experience of motor intentions, possibly in conjunction with earlier contributions from pre-SMA or SMA activity (4).

### M1 reflects intentions on a trial-by-trial basis

Next, we questioned whether finer-grain single neurons may track the subjective timing of W and A on a trial-by-trial basis. We restricted this analysis to the first day of recording to assure that a single units would not be included in duplicates. We examined sixty-six well-isolated units (examples shown in **Fig. 3A**).

**Figure 3.**
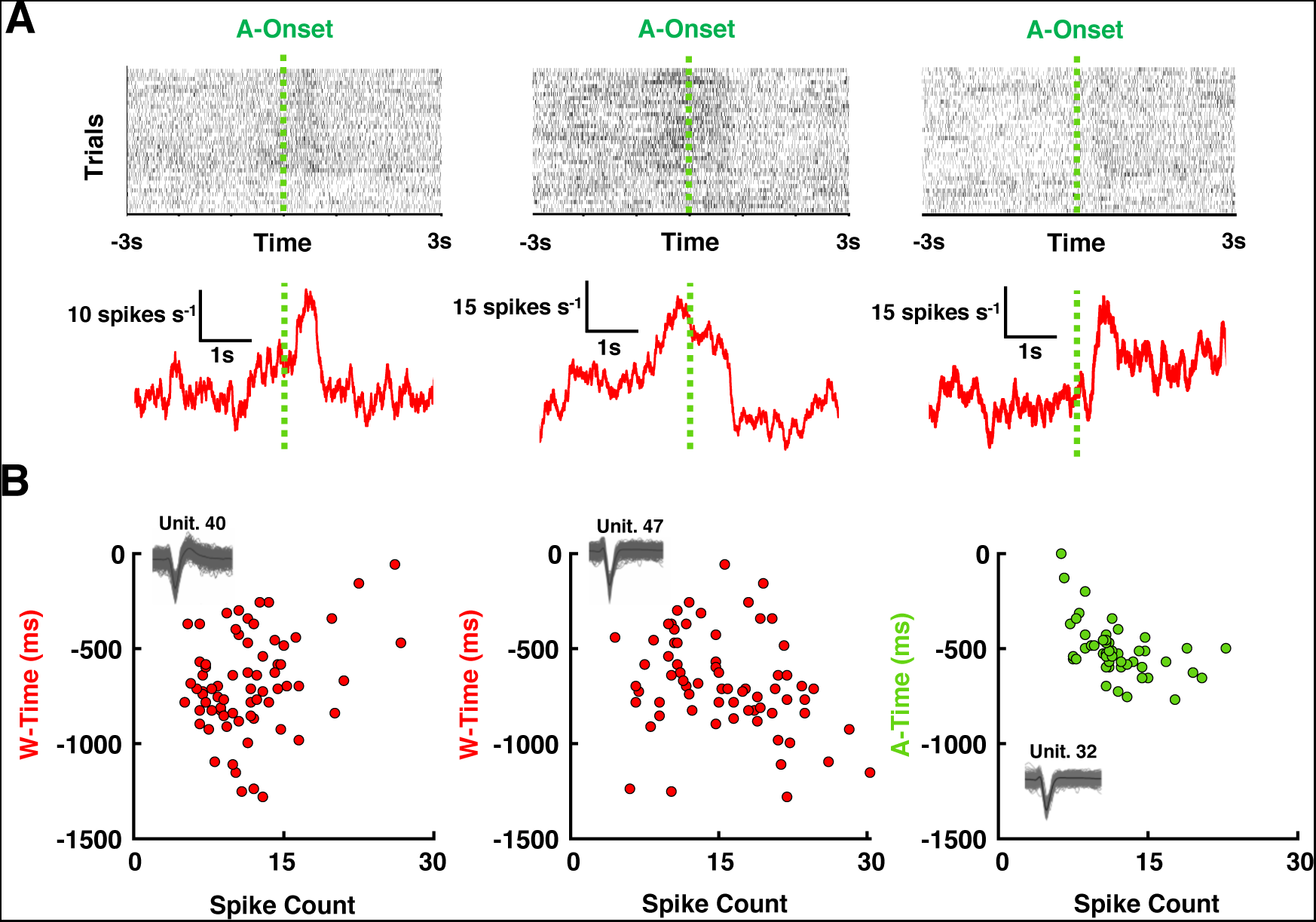
Single Unit Activity (SUA) in Primary Motor Cortex and Timing of Intentions and Actions. A) Raster Plots and average SUA of Example Units. Depiction of single trial activity of three representative single neurons. Top: The raster plots show every trial as a row and each black dot is a spike. Bottom: Evoked response surrounding time of action onset for the three representative neurons. **B) Correlation Between Spike Counts and Subjective Estimates**. Depiction of three representative correlations between the subjective timing of intentions (red, examples 1 & 2) and actions (green, example 3) relative to movement onset, and the spike-count of the neuron in the second prior to movement. No correlation is shown for the subjective timing of the tone, as none existed (8/66 for intention, 1/66 for action).

We performed a spike count during the 1000 ms-period preceding A. Eight neurons (12% out of 66) showed a significant correlation between their spike counts in this period and subjective W-time (all p<0.05 see **Fig. 3B, first** and **second column**). All 8 neurons are shown in **Fig. S6**. We note that Fried and colleagues’ (4) report ∼17% of neurons *responding* to W in frontal areas and ∼8% in temporal regions. Surprisingly, only a single neuron in the present study showed a correlation between its spike counts during this period and the subjective A-time (**Fig. 3B, third column,** r = −0.62, p = 0.01). We did not observe any neuron whose spike count correlated with E-time (see **Fig. S7** for similar results across different time periods).

Examining the spatial arrangement of the neurons showing a correlation with subjective W-time showed that all but one cell were on the anterior half of the electrode array (**Fig. S8**). This suggests a weak spatial arrangement, where M1 neurons closer to premotor cortex are more likely to relate to the perceived timing of intention (27).

### Real time decoding of intention

Lastly, we look at the impact of each of the elements of the intentional chain on the proficiency of real-time BMI decoding. First, we contrasted the decoder time-course during the full intentional chain to conditions missing a single element of the sequence. **Fig. 4A** shows that when actions were willed, there was a gradual rise toward the movement initiation threshold. Inducing a non-intended movement through NMES also affected the BMI classifier, but through a delayed and more abrupt rise toward threshold. These two conditions (i.e., intended vs. not) strongly bifurcated 1100ms prior to movement (p < 0.01; A-time at 0 ms), a timing that comfortably precedes the median timing of will perception during the full intentional chain (−597ms ± 79ms relative to A-time). When W did not lead to A, the decoder rose to threshold normally, but plummeted earlier (W-A-E vs. W-E differ 100ms post-movement onset, p < 0.01; **Fig. 4B**). Lastly, during the condition lacking E, the decoder was sustained for a longer period (**Fig. 4C**, WAE vs. WA differ 500 post-movement onset, p<0.01). These results suggests that the decoder (i) can index intention to move prior to the subjective experience (as is also highlighted in **Fig. 2F**) and (ii) that it is sensitive to all aspects of the intentional chain.

**Figure 4.**
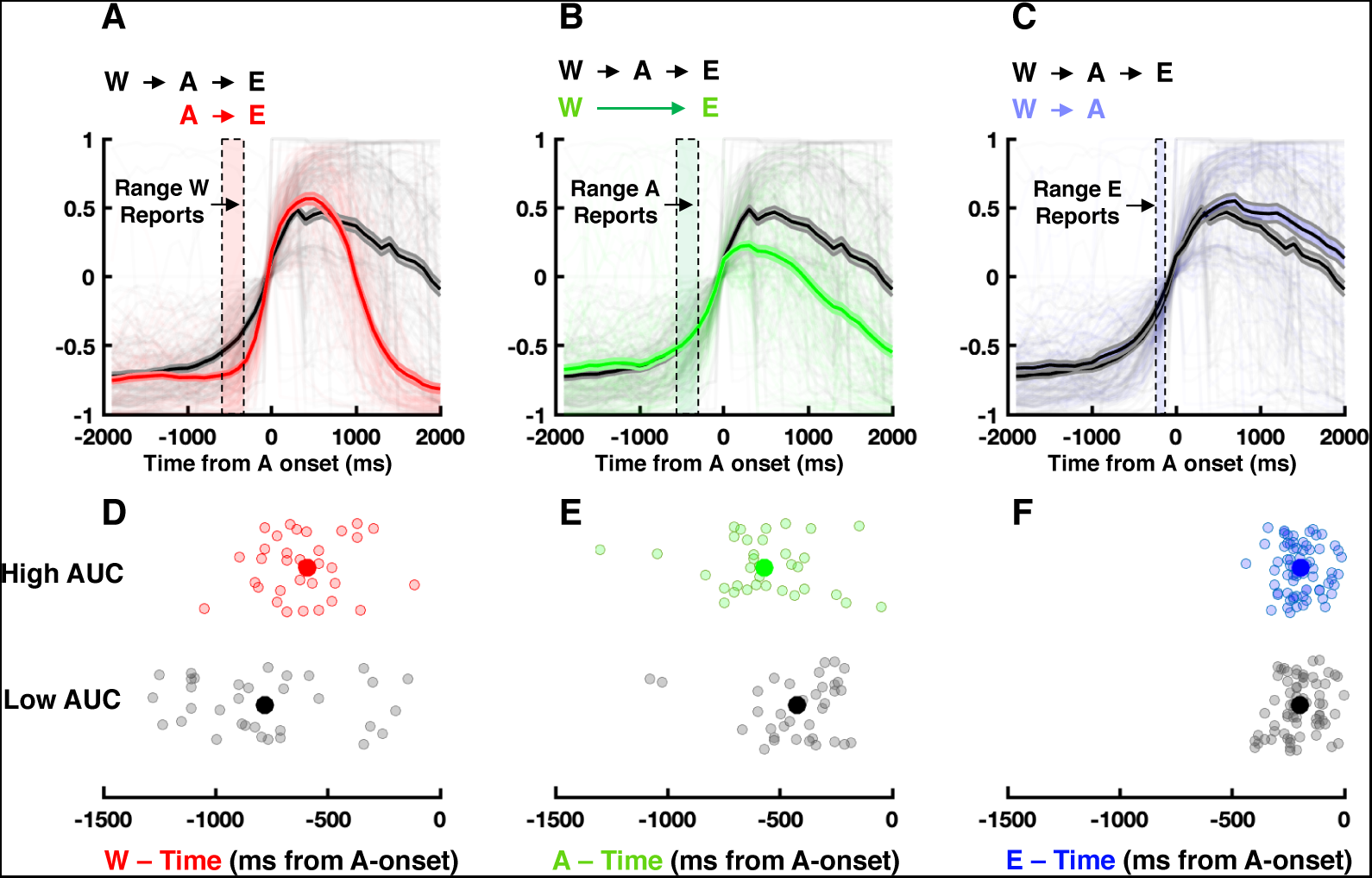
Movement Decoding Suggests that Deliberate Acts Bind Intention and Action. A) Effect of Intention. Contrast of the decoder time-course during the full intentional chain (black) and a chain without intention (red) suggests that intention causes an early and gradual rise toward the movement threshold (y=0). Shaded vertical area enclosed by dashed lines is the median ± 95%CI of W reports. **B) Effect of Action.** Contrast of the decoder time-course during the full intentional chain (black) and a chain without action (green) suggests that the decoder output decreases shortly after crossing movement threshold when no motor output occurs. Shaded vertical area enclosed by dashed lines is the median ± 95%CI of A reports. **C) Effect of Tone.** Contrast of the decoder time-course during the full intentional chain (black) and a chain without a consequence to movement (blue) suggests that the absence of tone results in a sustained heightened decoder output. Shaded vertical area enclosed by dashed lines is the median ± 95%CI of E reports. Shaded area around the decoder output is ± 1 S.E.M. All trials are shown in transparent in the background. **D) Decoder AUC and Intention.** Trials where the area under the decoder curve (AUC) was high (red) resulted in the perceived timing of intention to occur later (closer to action) in contrast to trials with a low AUC (black). **E) Decoder AUC and Action.** Trials where the AUC was high (green) resulted in the perceived timing of actions to occur earlier (closer to will) in contrast to trials with a low AUC (black). **F) Decoder AUC and Effect.** Trials where the AUC was high (blue) vs. low (black) resulted in similar perceived timing of the tone.

A final pertinent user-experience question is whether real-time BMI decoding reflects aspects of human intentional behavior. More precisely, we questioned whether performance of the decoder – indexed by the area under its curve (AUC) – may reflect in real time the unfolding phenomenology of intentional action. We coalesced all trials of a particular judgment type (i.e., W- time, A-time, or E-time). Then, we computed the AUC of the decoder for each trial and kept the top (high) and bottom (low) third of trials defined by AUC. Examining the participant’s reports – sorted by AUC of the decoder and independently from behavior – showed that in trials with high AUC, W was perceived as occurring later than on trials with low AUC (**Fig. 2D**, p = 3.77×10^-4^). Applying the same approach to A-time judgements, we found the opposite pattern: actions were perceived as occurring earlier (**Fig. 2E**, p = 0.01) in high versus low AUC trials. In other words, in trials where the BMI decoder was more proficient there was mutual attractive pull between intentions and actions, as determined behaviorally in the full intentional chain. The perceived timing of E was not modulated by AUC of the decoder (**Fig. 2F**, p = 0.22, see ***Supplementary Results*** and **Fig. S9** for a further control).

Overall, the M1 decoder indexes motor intentions prior to the subjective experience and its proficiency reflects the degree of subjective intention to move, as indexed by the subjective timing of W and A. These findings show that a M1 BMI decoder can track the *subjective experience* of intention to move in (near) real time.

## Discussion

We leveraged the unique opportunity to selectively bypass intentions, actions, or the corollary consequences of these in a tetraplegic person fitted with an intracortical M1 implant and NMES leading to movements of his own body. Results demonstrated a novel form of intentional binding, not between actions and effects (11, 12), but between intentions and actions. Interestingly, the time distortions caused by intention-action binding are stronger than those from action-effect binding (see ***Supplementary Discussion***). Neurally, we demonstrate that unit and population-level dynamics index the presence of intention. Critically, having access to subjective timings, we also show that neural activity in M1 shows an evoked response corresponding to subjective onset of intention, and these covary on a trial-by-trial basis. Lastly, our data show that a neural decoder in M1 can track in near (∼100ms) real time the subjective degree of intentionality, as indexed by intention-action temporal binding.

Fried and colleagues (4) showed that neurons in the pre-SMA, SMA, and ACC all anticipate the subjective timing of intention by about 700-1500ms. Here, we close a critical gap in knowledge by recording from M1, an area downstream from those previously probed. We show that evoked spiking activity in M1 co-occurs with and does not precede (as in pre-SMA, SMA, and ACC) the experience of intention. This suggest that while (multivariate) signals related to intention may be present (i.e., are decodable) in M1 and elsewhere (9, 23–25, 28–33) prior to the onset of the subjective experience of intention, this latter one coincides with evoked activity in M1. This observation, and our emphasis on relating neural activity with the subjective onset of intentionality, breaks with the dominant view that ascribes intention at ever-earlier stages of the cortical action hierarchy (34–36). Together with previous non-invasive studies (4, 12, 37), our BMI-NMES data suggest that while other frontal and posterior parietal areas are likely involved in initiating the drive to move, M1 activity reflects the onset of the subjective experience of intention. These observations are acknowledgedly from a tetraplegic patient and an experienced BMI user who may have experienced a degree of neuroplasticity. Future work will therefore need to determine how these results relate to the general population. However, we highlight that true and independent control of intention and motor output, as developed in the current series of experiments, is only possible in a BMI-NMES context.

From a neuro-engineering perspective, the decoding results provide direct evidence that M1 decoders are not only built to translate motor intentions into movements, but that they can index motor intentions prior to their associated subjective experience. Moreover, we leveraged our new implicit measure of intention – the subjective temporal attraction between intention and action (W-A binding) – to further ascertain the relation between real-time motor decoding in M1 and the degree of subjective intentionality. We show that the greater the proficiency of decoding the user’s intention to move, the stronger is the subjective W-A binding. This provides further evidence for M1 as key node in the cortical intention network, and demonstrates that M1 activity encodes the degree of subjective intentionality. This has therapeutic impact and suggests that intentional signals ought to be leveraged in future generation BMIs.

Our findings add scientific insight into the timely ethical and philosophical debate regarding the broader impact of novel neurotechnologies on individuals and societies. In a not-so-distant future, neurotechnologies may not only sense an individual’s intentions before they manifest in the environment, but may also allow outsiders to monitor the degrees of intentions that an agent may experience. This underscores the urgent need to better understand the neural correlates of our sense of free will and agency (38–40), protecting people’s mental privacy and integrity, and fostering new neurotechnological therapies.

## Methods

### Participant

The subject was a 27-year-old male with stable, non-spastic C5/C6 quadriplegia from a cervical spinal cord injury (SCI) sustained 8 years prior to the current experiment. The participant’s International Standards for Neurological Classification of SCI neurologic level is C5 (motor complete) with a zone of partial preservation to C6 (41). He is an expert BMI user, with over 5 years of usage (23). The subject has full range of motion in bilateral shoulders, full bilateral elbow flexion, a twitch of wrist extension, and no motor function below the level of C6. His sensory level is C5 on the right (due to altered but present light touch on his thumb) and C6 on the left. He has intact proprioception in the right upper limb and the shoulder for internal rotation through external rotation, at the forearm for pronation through supination, and at the wrist for flexion through extension. Proprioception for right digit flexion through extension at the metacarpal-phalangeal joints was impaired for all digits. Approval for this study was obtained from the US Food and Drug Administration (Investigational Device Exemption) and The Ohio State University Medical Center Institutional Review Board (Columbus, Ohio). The participant referenced in this work completed an informed consent process before taking part in the study. He also provided written permission for photographs and video.

### Surgical Procedures and Data Acquisition

The patient underwent a left frontoparietal craniotomy for implantation of a Utah microelectrode array (Blackrock Microsystems Inc, Salt Lake City, Utah) in the (dominant) hand region of primary motor cortex. This region was identified via pre-operative functional Magnetic Resonance Imaging where the patient was asked to imagine performing hand movements (**Fig. 1**). The area was targeted via an intraoperative navigation system (see (23) for details). The array contained 96 channels (4.4mm x 4.2mm x 1.5mm in depth) and was implanted into the cortex using a pneumatic inserter. Reference wires were placed subdurally. Neural data was sampled at 30kHz and hardware band- pass filtered between 0.3Hz and 7.5kHz (3^rd^ order Butterworth). Once digitized, the data were transmitted to MATLAB (Mathworks, Natick, MA) in 100ms bins, where signal artifacts due to NMES were removed by blanking over 3.5ms around the artifact (defined as signal amplitude >500μV simultaneously in at least 4 of 12 randomly selected channels).

### Movement Decoder

A non-linear Support Vector Machine (SMV) was used to translate neural activity to intended hand movements. The inputs to the SVM were mean wavelet power (MWP) over 100 ms bins. That is, neural activity was decomposed into 11 wavelet scales (Daubechies wavelet, MATLAB), and the coefficient of wavelets 3-6, corresponding to the multi-unit frequency band spanning from 235 to 3.75kHz, were then normalized and averaged within 100 ms bins resulting in 96 values, one for each channel. Training data for the decoder was obtained by prompting the participant at the beginning of each recording session to imagine performing a specific hand movement (HO or HC) using an animated virtual hand displayed on a computer monitor. The subjects performed 7 blocks of classifier training (5 repetitions per hand movement type) prior to starting the main experiments (**Fig. S1**). Training took approximately 10 minutes per session. Output classes were built for each movement and had scores that ranged from −1 to 1. The appropriate Neuromuscular Electrical Stimulation (NMES, see below) became active when the output score for a given movement exceeded zero.

### Neuromuscular Electrical Stimulation

A Neuromuscular Electrical Stimulation (NMES) system was used to evoke hand movement by stimulating forearm muscles. The NMES system consisted of a multi-channel stimulator and a flexible, 130-electrode, circumferential forearm sleeve. The coated copper electrodes were 12mm in diameter, spaced at regular intervals in an array (22mm longitudinally x 15mm transversely), and delivered current in monophasic, rectangular pulses at 50Hz (pulse width 500μs, amplitude 0- 20mA). Desired hand movements were calibrated at the beginning of each session by determining/confirming the intensity and pattern of electrodes required to stimulate intended movements. This procedure took approximately 5-10 minutes per session.

### Experimental Design

Our goal was to establish the behavioral and neural ramifications of breaking the normal course of events between intentions to move, movement, and the consequence of some movements. In turn, from a baseline rest position, the subject was asked to perform two movements, right hand opening (HO) and closing (HC) – i.e., the decoder was trained to recognize these two neural patterns. Hand closing resulted in an auditory tone being played 300 ms after movement onset (or the decoder surpassing movement threshold). The analyses in the current report focus on H.C. trials, as these were associated with consequences in the external environment (i.e., the tone), and thus had all components of a full intentional chain. While performing these movements, the subject observed the rendering of a clock with a single hand that would complete a full cycle in 2560 ms (5). The initial position of the clock hand was random on each trial. Over the course of 12 separate sessions (∼ 2-4 hours/session including setup) the subjects completed 5 different experiments, containing variations where he reported either the time he willed to move (W-time), the time of action onset (A-time), or the time of sound onset, the effect of movement (E-time).

In the first experiment, on different blocks we randomly activated the NMES resulting in H.C., or we presented the tone at a random time without NMES (∼40-50 repetitions per condition). This experiment was a baseline experiment to assess the subject’s temporal perception of movement and auditory stimuli in the absence of any causal structure (i.e., no intention to move, nor a movement causing the effect in the external environment). The second experiment was the “full intentional chain” where intention resulted in movement, and a particular movement (H.C., but not H.O.) resulted in a tone being played. On different trials the subject reported the timing either of his intention, movement onset, or tone onset (∼40-50 repetitions per condition). The third experiment broke the expectation that H.C. result in external consequence, and on different trials the subject was asked to report the timing of his will to move or actual movement (∼40-50 repetitions per condition). In the fourth experiment the subject was instructed not to imagine moving, and instead at a random time we activated the NMES resulting in H.C. Thus, in this experiment there was movement and an effect to this movement, but no intention. On different trials the subjects indicated either movement or tone onset (∼40-50 repetitions per condition). Lastly, and quite uniquely given the experimental setup (BMI coupled with NMES), in the fifth experiment the subject willed to move, and this resulted in the decoder crossing the threshold that would ultimately evoke a consequence, the tone, but no movement. On different trials we asked the subject to report either when he first intended to move, or the time of sound onset (∼40-50 repetitions per condition). Report type (W, A, or E) was varied in “mini-blocks” of 5 trials, and a full experiment was not run in 1 session, but instead experiments were intermixed between sessions.

### Analyses

Behavioral estimates of each judgment type (W, A, E) and different combinations of elements of the intentional chain (**Fig. 1B**, different rows) were coalesced and their circular median computed. Targeted non-parametric statistics were conducted.

Regarding single-unit spiking activity, data files were concatenated across an entire session, filtered with a 300Hz-30kHz bandpass filter, and spike sorted via Wave_Clust in MATLAB (42). Only data from 1 session was used in the analyses for single unit spiking activity, to prevent “double dipping” (i.e., counting repeat observations as independent). Clusters were then manually inspected for waveform shape and violation of the inter-spike interval. Spikes were binned in 1ms intervals and then spike counts were performed. Correlations with subjective timing of intention, actions, and effects were computed by Pearson correlation. For firing rate plots in **Fig. 3A**, binned spikes were convolved with a Gaussian distribution with a standard deviation of 50ms.

Multi-unit spiking activity from all 96 recording channels were separated as a function of judgment type and the different combinations of the intentional chain. Spikes were then convolved with a Gaussian distribution (50ms standard deviation). We summarized this activity by averaging each channel across trials. The onset of evoked MUA was defined as the signal being 5 times the noise level.

To account for spiking activity of the population of neurons as a whole, we performed a principal component analysis (PCA) on the average firing rate for each condition. We kept the two components that explained most variance. For statistical analyses we re-sampled trials (with replacement) 1000 times, before conducting PCA, and determined the 95% confidence intervals.

Lastly, regarding the neural decoder analyses, the outputs of the decoder were amalgamated as a function of judgment type and different combinations of the intentional chain. Statistical contrasts as specified in the main text (e.g., WAE vs. AE) were conducted via time-resolved unpaired t-test.

## Acknowledgements

The authors thank Ian for his dedication to the study and insightful conversations. AS is supported by the Swiss National Science Foundation (grant PP00P3_163951/1), OB is supported by the Swiss National Science Foundation and the Bertarelli Foundation. MB was supported by the Craig H. Neilsen Foundation (Grant number: 651289) and State of Ohio Research Incentive Third Frontier Fund. JPN is supported by K99NS128075. The funders had no role in study design, data collection and analysis, decision to publish or preparation of the manuscript.

## Author Contributions

J.P.N., O.B., and A.S designed the experiment. M.B., S.C. A.R. implemented the experimental set-up. J.P.N., M.B., S.C., M.S., P.O. collected the data. J.P.N. analyzed the data. J.P.N. and A.S. wrote the manuscript. All authors edited the manuscript.

## Supplementary Text

### Supplementary Results

Behavior. The presence of E did not modulate the perceived timing of W and A. At first glance, this seems contrary to previous reports of intentional binding effects (11, 12). Thus, we also asked the BMI participant to report the timing of A (**Fig. 1B, Row V**) and E (**Fig. 1B, Row VI**) in the absence of other elements of the intentional chain. We compared these estimates with control events in which unintended actions were followed by similar effects (**Fig. 1B, Row II**). This showed that A was perceived as earlier (p = 0.01) and E are perceived as later (p = 2.2 x 10^-4^) when each event occurred in isolation, as compared to when they occurred in association with each other. This pattern of response reproduces the classic temporal binding effect, but also extends previous reports suggesting that a causal, and not necessarily an intentional relation between actions and effects may be sufficient to engender a degree of attractive pull between one another (43–46). Nevertheless, the fact that some binding may occur in the absence of intention does not preclude the possibility that an additional, and specific element of binding occurs when intention is additionally present (47).

Neural Decoding. As a control, we demonstrate that the AUC of the decoder did not differentiate between the objective timing of action relative to trial onset (as opposed to A-onset in the **Main Text**), for neither trials where the BMI participant was asked to estimate the timing of W, A, or E (all p > 0.17. **Fig. S9**).

### Supplementary Discussion

Behavior. The timing judgments revealed a strong and previously unmeasured perceived temporal binding between intentions and actions. This resembles the previously described temporal attraction between actions and effects described in the classic intentional paradigm (11, 12). In fact, in the present BMI-NMES context we also observed, albeit to a lesser extent, binding between actions and effects. This action-effect binding has traditionally been considered to reflect a functional need to bind together sensorimotor events that surround voluntary actions, and that this function may be critical for the normal experience of agency. Temporal binding has now been described in several other experimental contexts, including those without motor intentions. Based on these recent data, several groups (43–46) have suggested that the traditional intentional binding may not be driven by intentional processes per se, but by inferred causality. That is, the repeated occurrence of a specific effect after a specific action may bind the two by multisensory “causal binding” (see (48), for a demonstration of temporal binding as cue integration and **Fig. 1B**, **Row II** vs. **Rows V-VI** for corroborative evidence). In turn, the most parsimonious explanation of the intention-action binding herein described may be perceived causality. Moreover, we argue that the stronger temporal binding between intention and action, rather than between action and effect may reflect the lifelong association between willing to move and the resulting movement of the body. That is, there is generally an unequivocal association between intending to move and movement, while the binding between an action and a given effect in the external world depends on the contingent and variable causal structure of the environment. Second, it may be argued that the neuro-cognitive resources demanded by action initiation – particularly in the present BMI-NMES context – outweigh those involved in perceiving the environmental effect and thus may contribute to the greater binding of action toward intention rather than toward the effect (see (19) for a similar argument and empirical evidence). Together, the present behavioral findings demonstrate a new and stronger than the classic (11, 12) form of intentional binding between intention and action.

### Supplementary Figures

**Figure S1.**
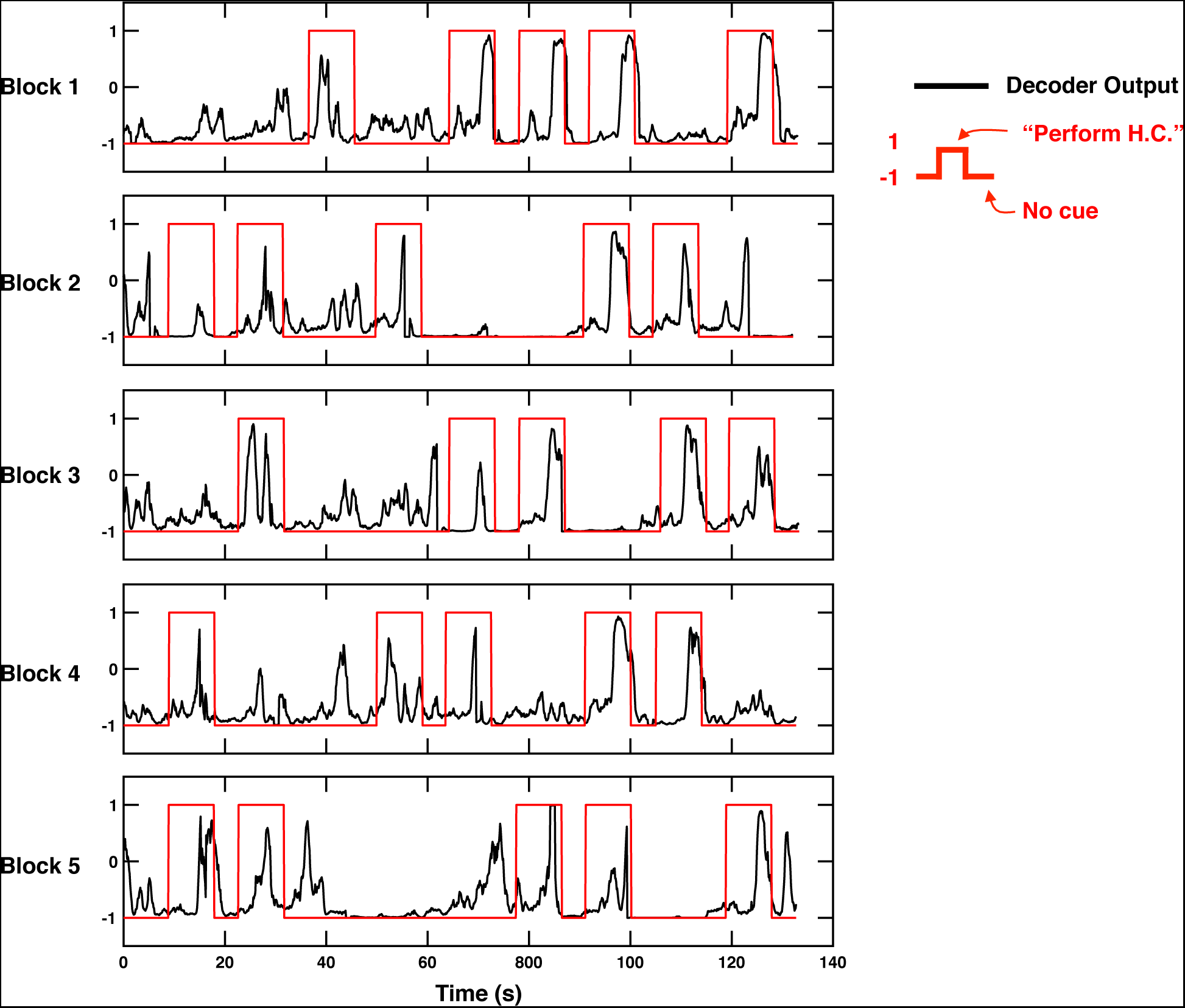
Evoking Hand Closure via M1 Decoder. The BMI user was asked to perform H.C. movements in a self-paced manner, but only during specific intervals (each internal being ∼11 seconds long). These intervals are marked by the red curve being equal to 1. The decoder output spanned between −1 and 1, and would cause movement via NMES after zero crossing. Trials were performed in short blocks of 5 trials, and the figure shows 5 example blocks of 5 trials. Decoder output is represented in black. As indicated in the main text, during the course of the entire experiment (12 sessions of ∼3-4 hours/session), the user was able to produce H.C. on 89.2% of requests.

**Figure S2.**
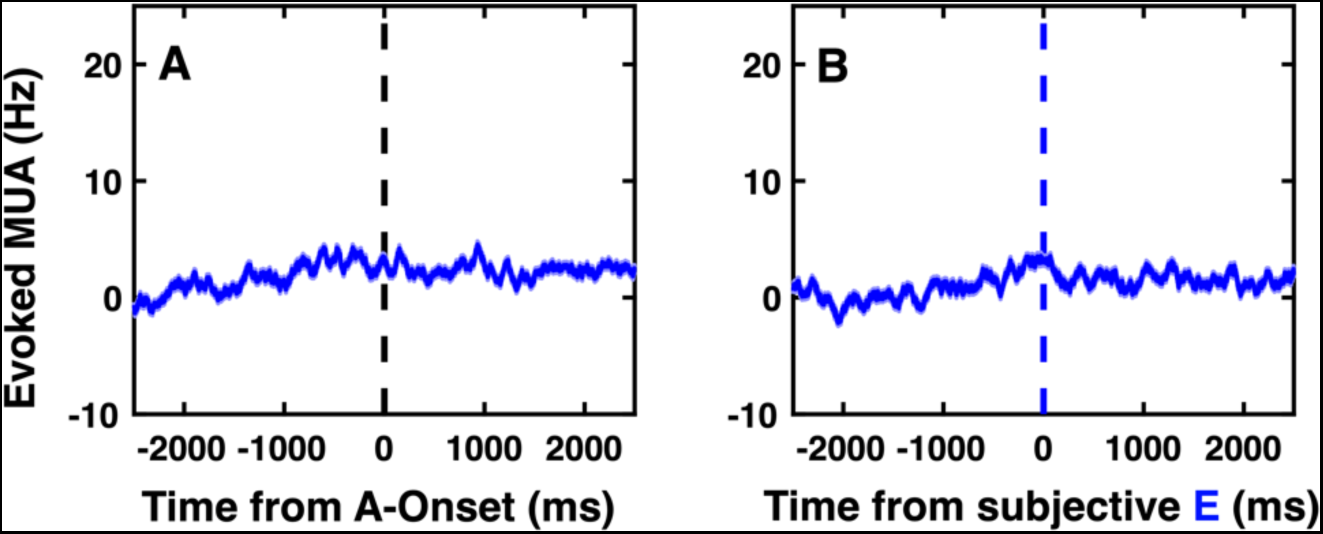
Evoked Multi-Unit Activity by E. **(A)** MUA evoked by the tone, aligned to movement onset. (**B)** MUA evoked by the tone, aligned to the subjective timing of the tone occuring. Shaded areas surrounding the average MUA are S.E.M. No evoked response is observed.

**Figure S3.**
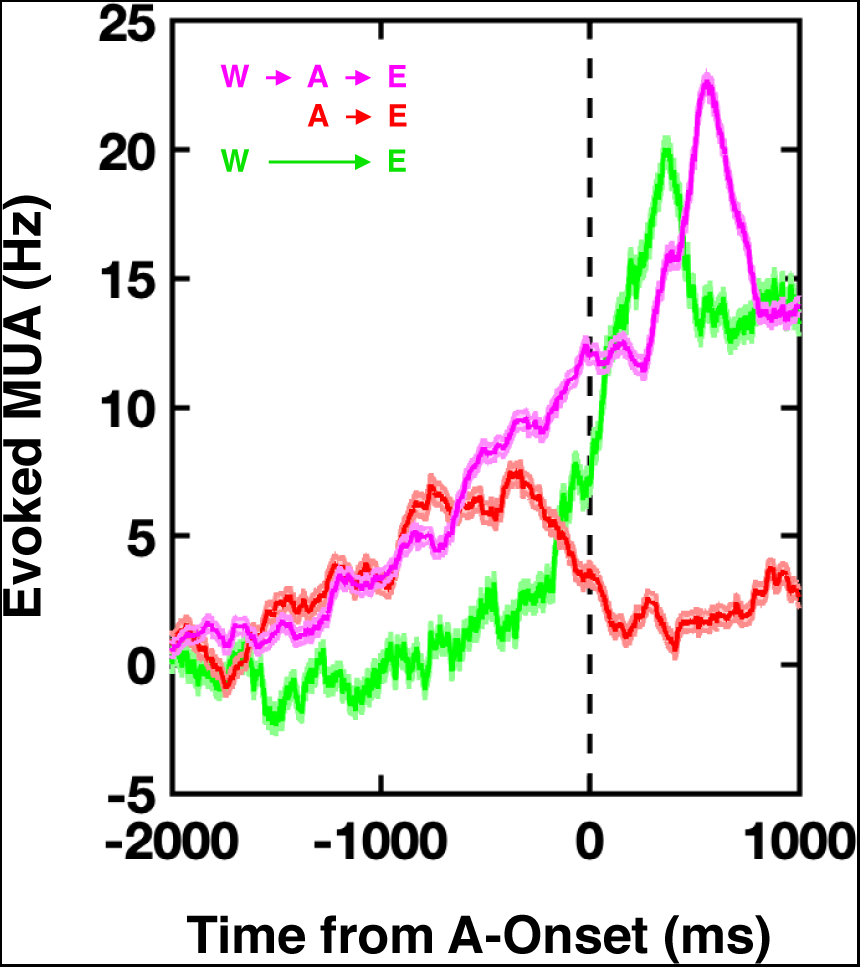
Evoked Multi-Unit Activity (MUA) during the full intentional chain (WAE, purple), as well as during cases missing W (red) or A (green). MUA is aligned to the onset of action, or in the case of no-action trials, to when the decoder reached threshold that would have evoked an action. Shaded areas surrounding the average MUA are S.E.M. Of note, when actions are intended, there is a gradual increase in firing rate prior to movement onset.

**Figure S4.**
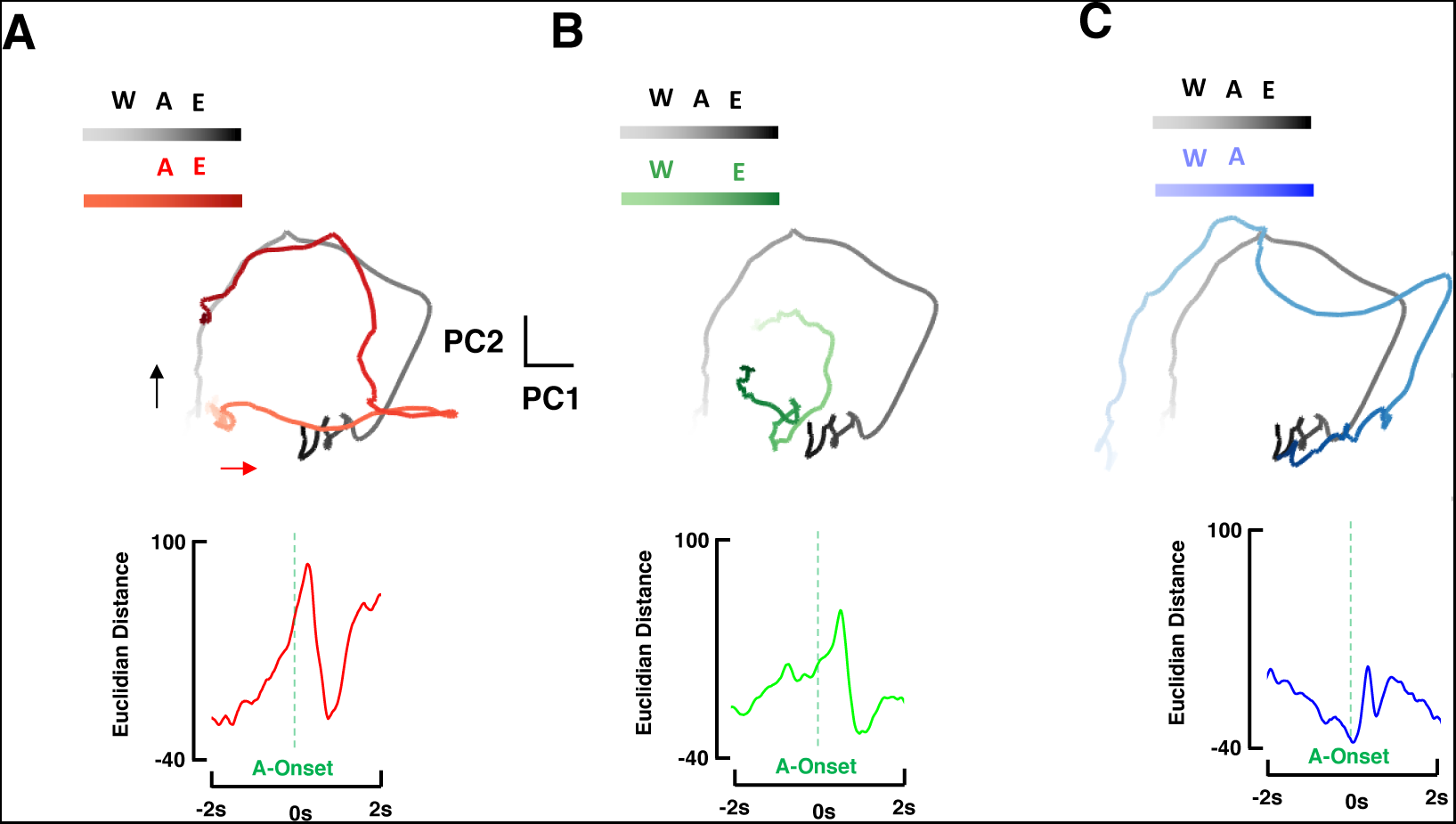
Population-level Dynamics in Primary Motor Cortex and Presence of Intentions, Actions, and Effects. **(A).** Top: Population dynamics on trials with the full intentional chain (black) and trials solely missing W (red). The two principal components accounting for most of the spiking variance (multi-unit activity) are plotted. Overall these components accounted for 81.9% of multi-unit activity and their shape in latent space showed the circular pattern stereotypical of M1 (Churchland et al., 2012). Hue contrast increases with time. Bottom: Euclidian distance as a function of time from movement onset. The distance in latent space between neural trajectories with or without intention, as a function of time from movement onset. (**B)** and **(C)** follow **(A)**, but for trials with and without action (B, green) and effect (C, blue).

**Figure S5.**
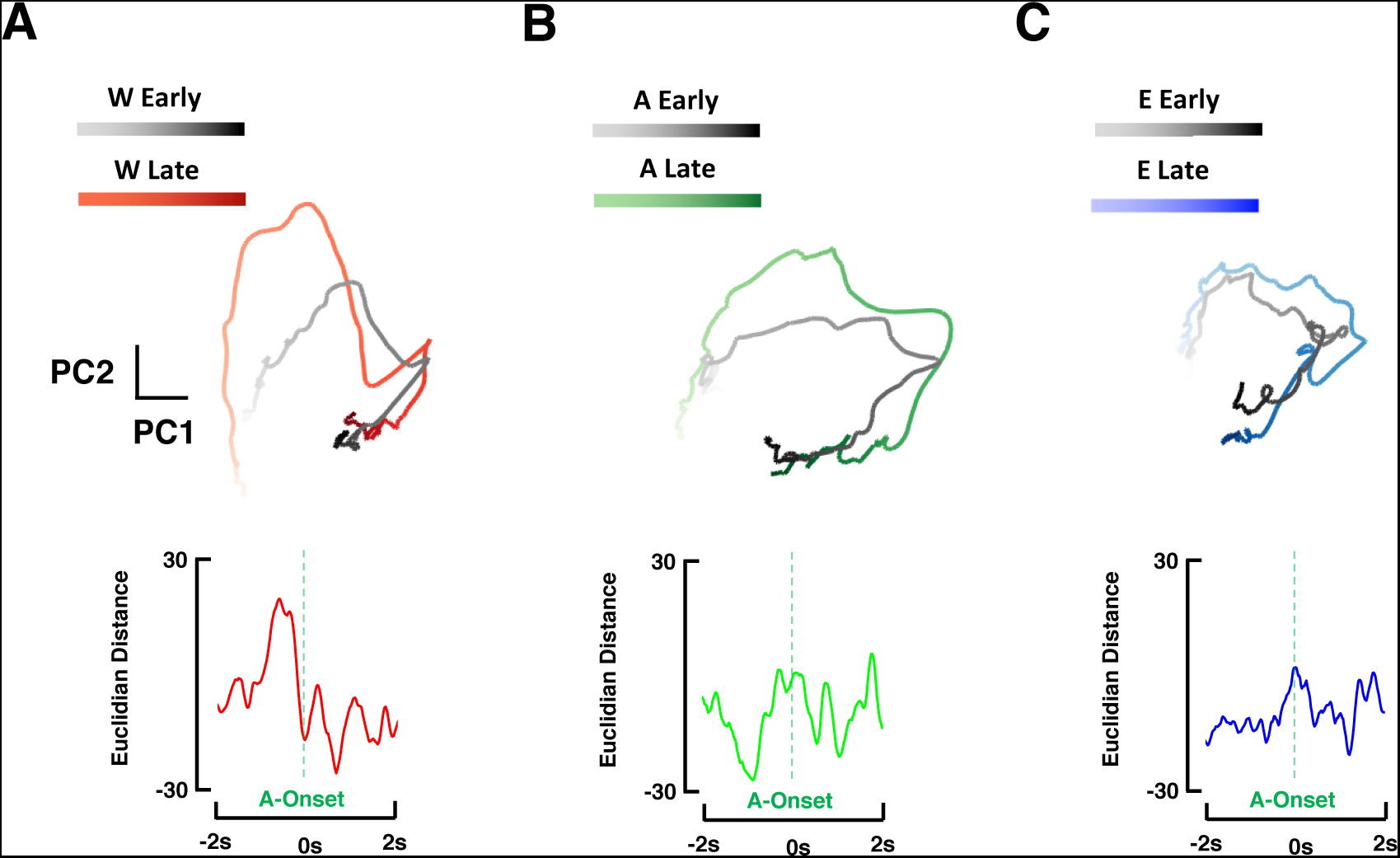
Population-level Dynamics in Primary Motor Cortex and the Relative Subjective Timing of Intentions, Actions, and Effects. **(A).** Top: Latent trajectories of trials with intention, perceived relatively early (black) or late (red). Bottom: Euclidian distance between these trajectories, as a function of time from movement onset. **(B)** and **(C)**, are as **(A)**, but separating trials as a function of the subjective timing of actions (B, green) and effect (C, blue).

**Figure S6.**
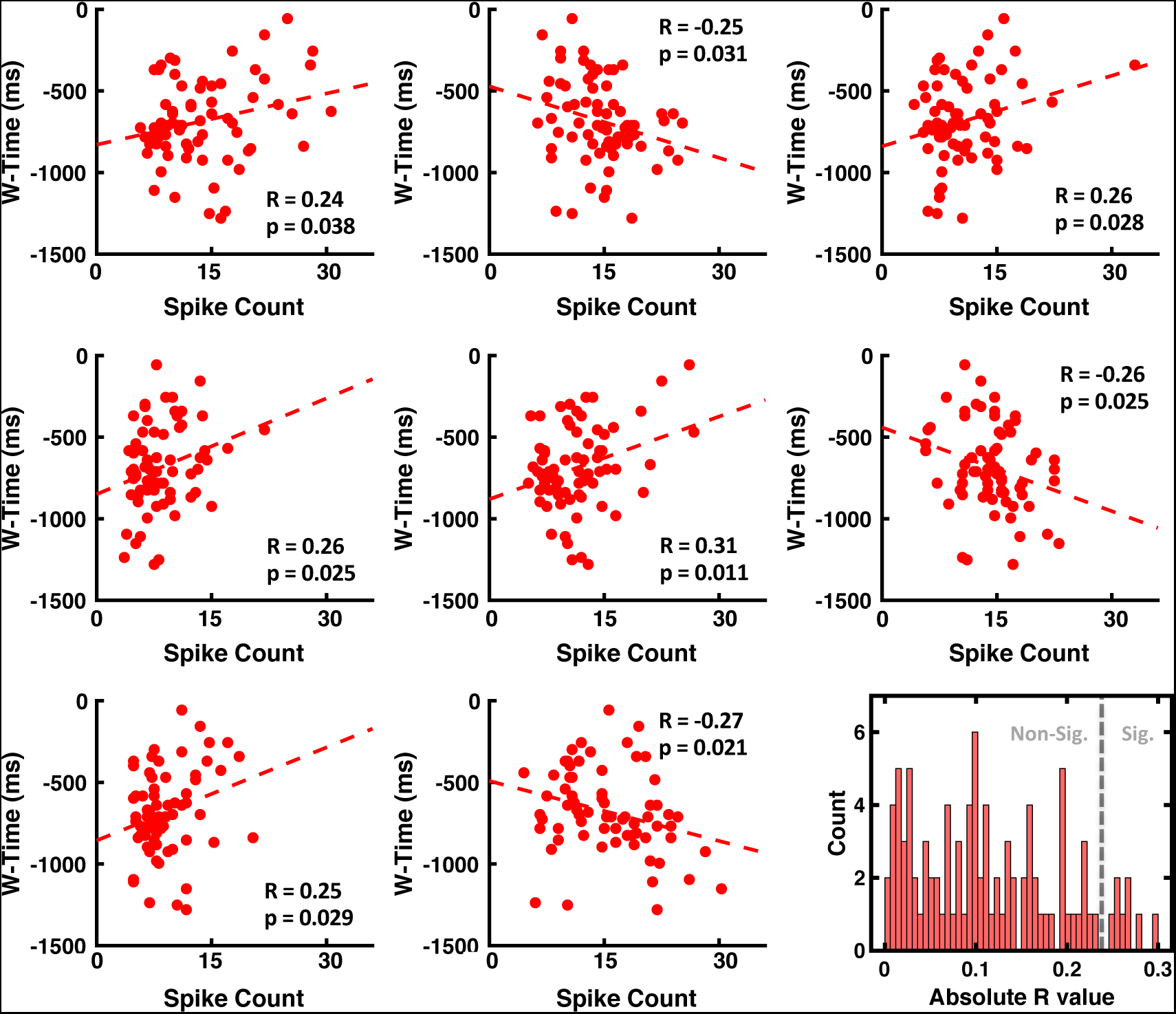
Correlation between spikes counts in the second prior to movement and reported time of intention relative to movement. The first eight panels show correlations between spike counts in the second prior to movement (x-axis), and the reported time of intention (y-axis, W-time). The last panel, lower right, shows the distribution of the absolute r-values.

**Figure S7.**
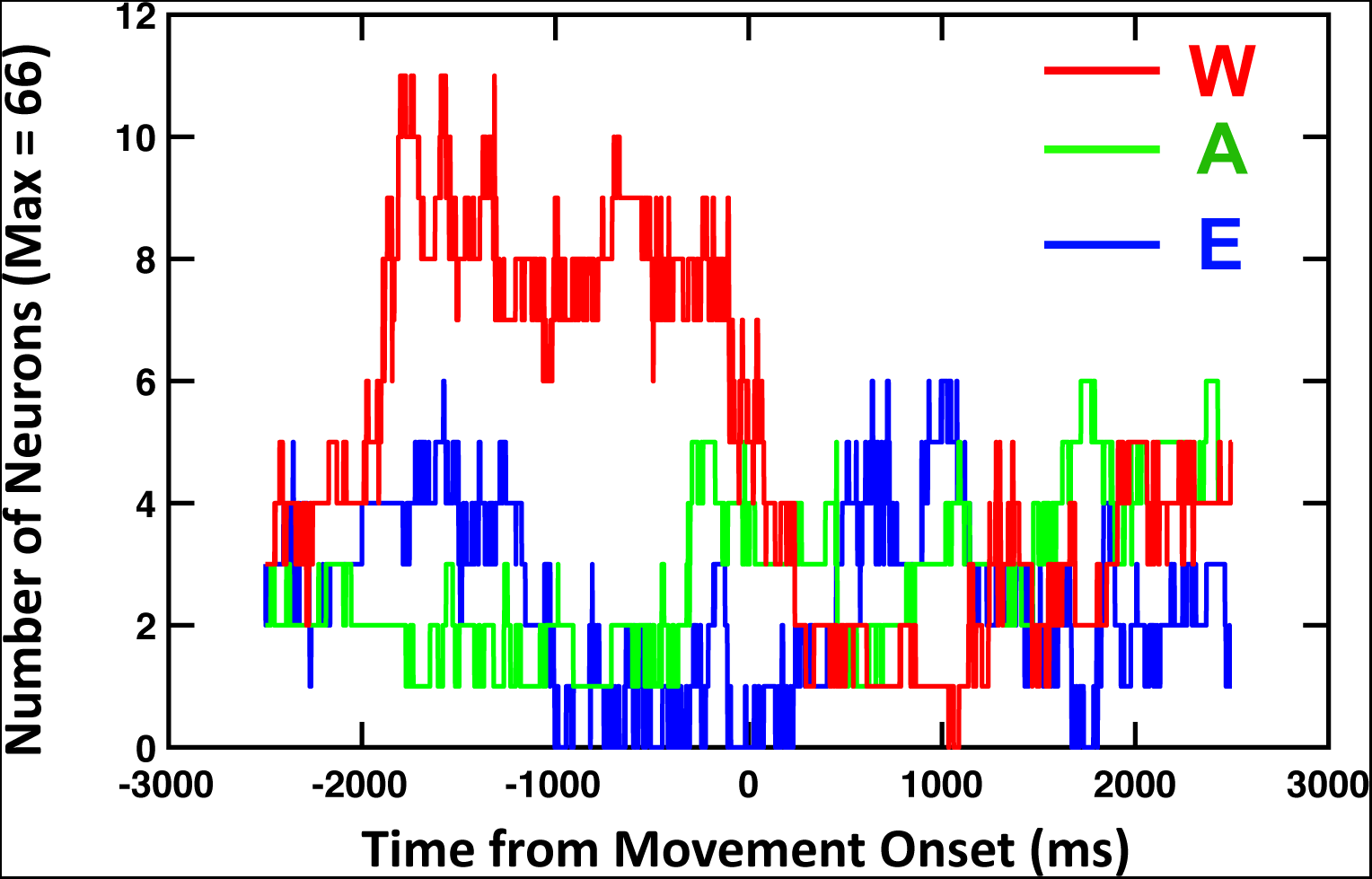
Number of Neurons Correlating with Estimates of Intention (W), Action (A), and Effect (E). Spike counts were performed within a window of 1000ms, and correlated with the subjective estimate of the timing of intentions (red), actions (green), and the corollary effect of the action (tone, blue). The spike-count time window was moved 1ms at a time, and the number of neurons demonstrating a significant correlation (p<0.05) is plotted as a function of time (centered in the middle of the time-window; e.g., x = −1000ms, means the time-window was from −1500ms to −500ms with respect to movement onset). Results demonstrate a clear over-representation of neurons correlating with the subjective timing of intentions rather than actions or effects. This effect is sustained from ∼ −2000ms to 0ms post-movement onset. After the movement, no estimate (intention, action, or effect) more frequently correlated with single-unit spiking activity.

**Figure S8.**
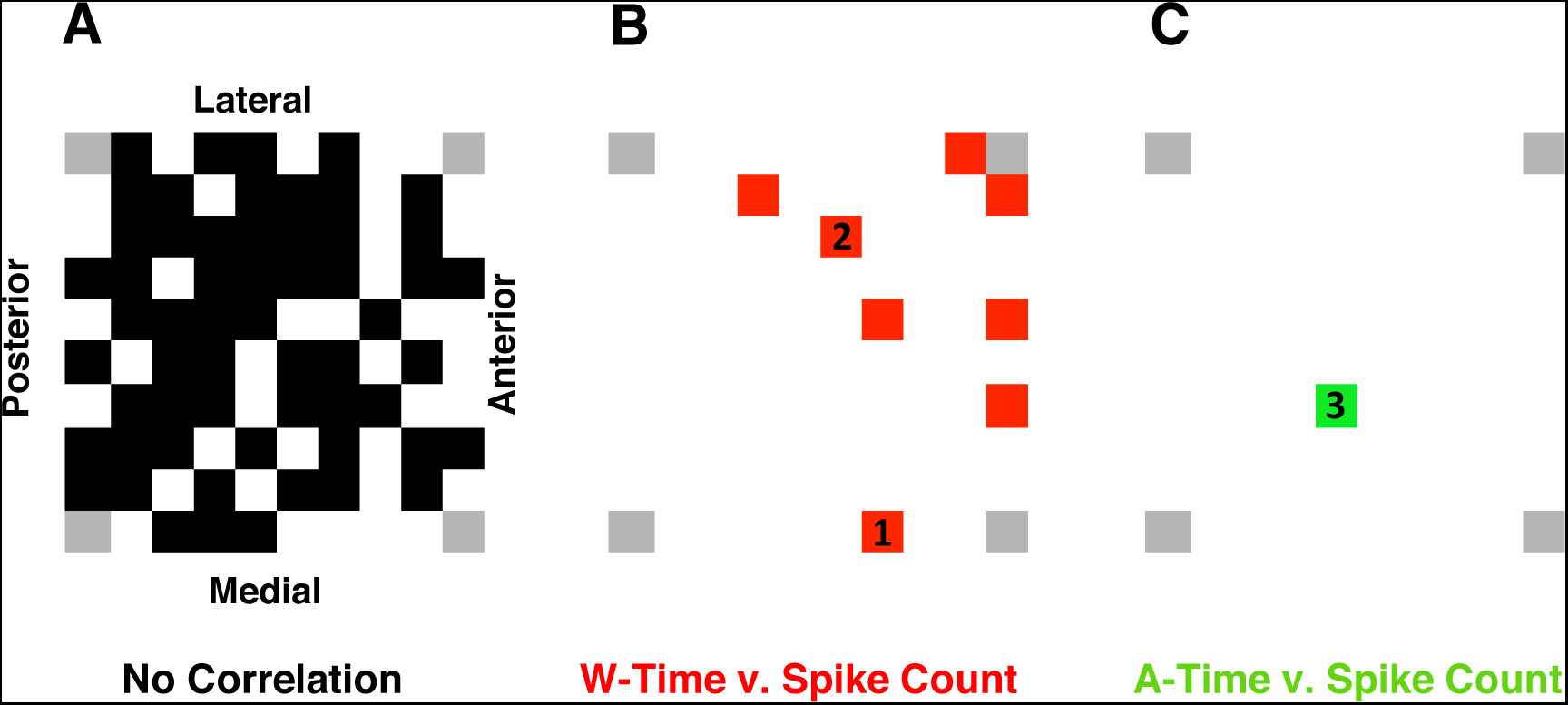
Topographical Arrangement of Electrodes where Correlation Exist between Spiking Activity and Timing Estimates on a Trial-by-Trial Basis. **(A)** In black, location on the Utah array where neurons were recorded but showed no correlation with timing estimates. The anterior portion of the array (facing pre-motor cortex) is on the right. **(B** In red, location of electrodes recording a neuron showing a correlation with the subjective timing of intentions. **(C)** In green, location of the electrode recording the neuron showing a correlation with the subjective timing of action. In **(B)** and **(C)** locations demarked (1), (2), and (3) correspond to neurons shown in main text, Figure 3B.

**Figure S9.**
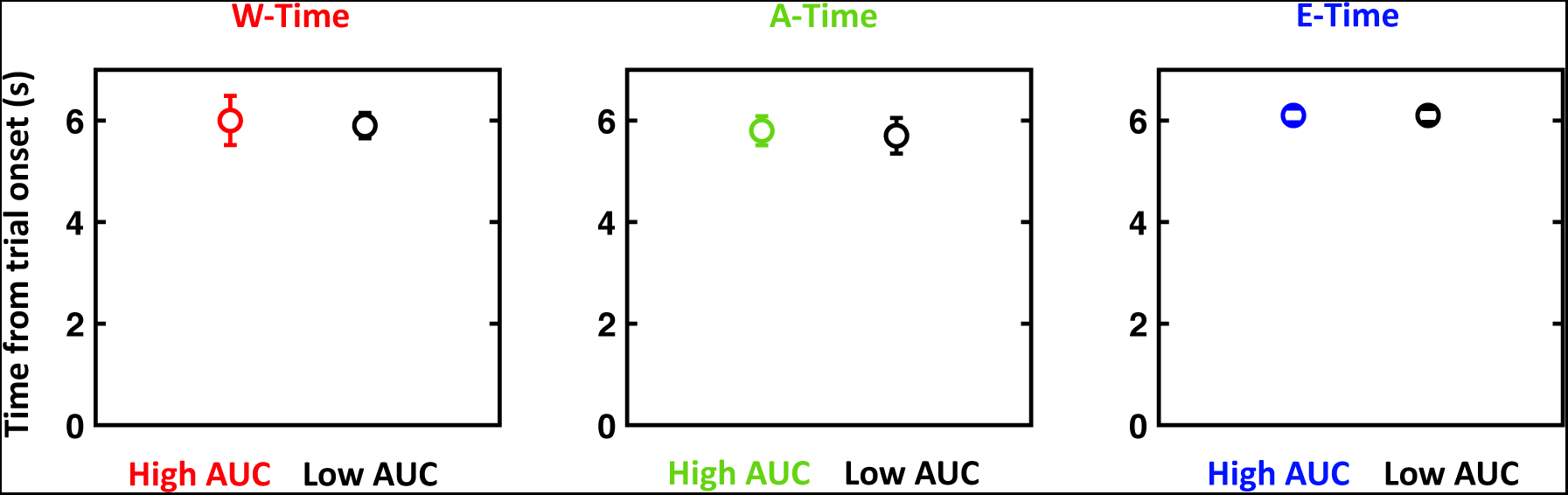
Time to Movement Onset as a Function of Decoder Area Under the Curve. Trials were split as a function of their total AUC of the movement decoder and estimate type – timing of intention (red), movement (green), and tone (blue). The AUC did not discriminate for how long it took the decoder to cross the threshold for movement relative to trial onset.

